## Supplementary figures and images for "Opposing, spatially-determined epigenetic forces impose restrictions on stochastic olfactory receptor choice"

### Supplemental Figure 1

Supplementary Figure S1

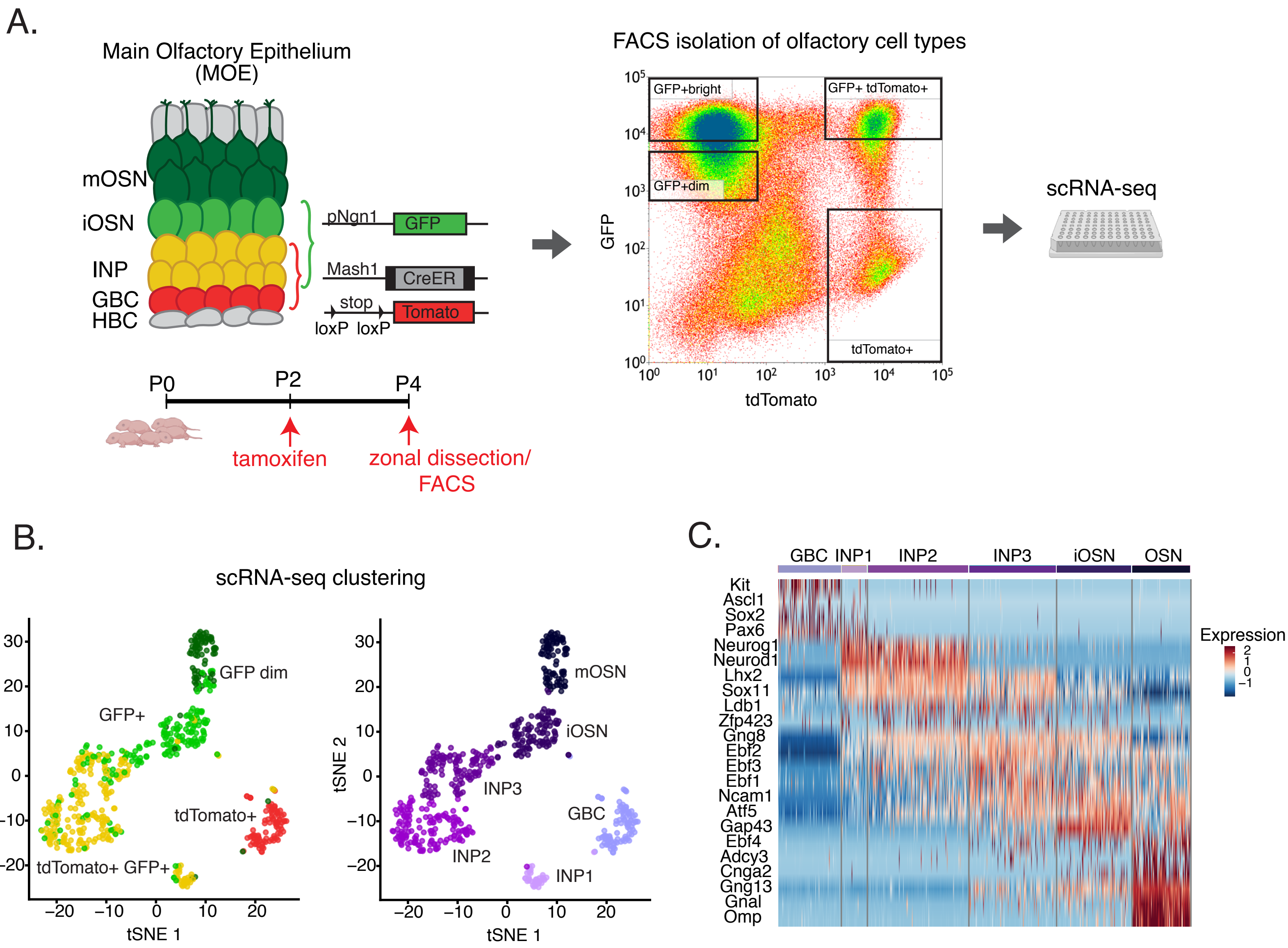

### Supplemental Figure 2

#### Supplementary Figure S2

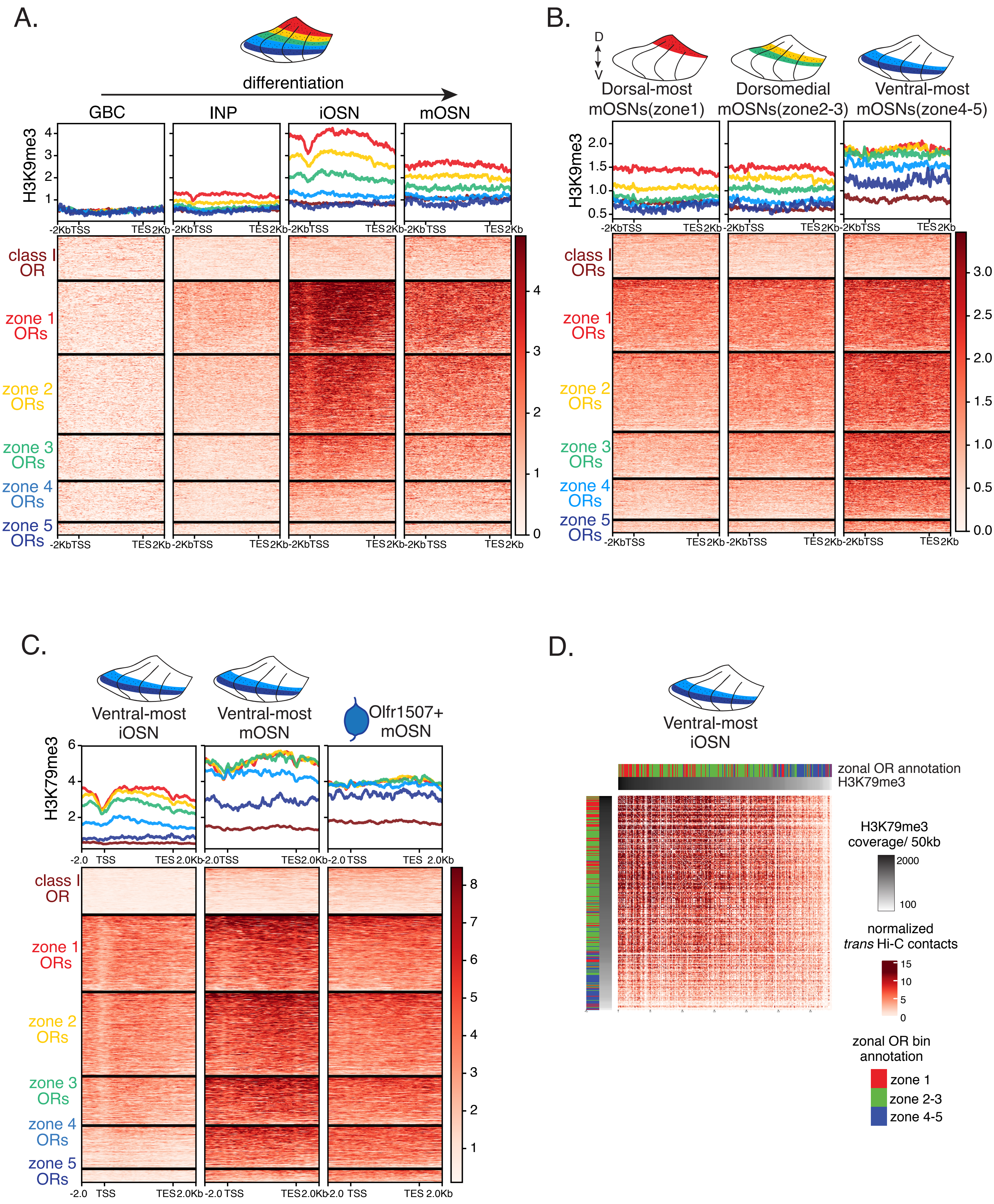

### Supplemental Figure 3

Supplementary Figure 3

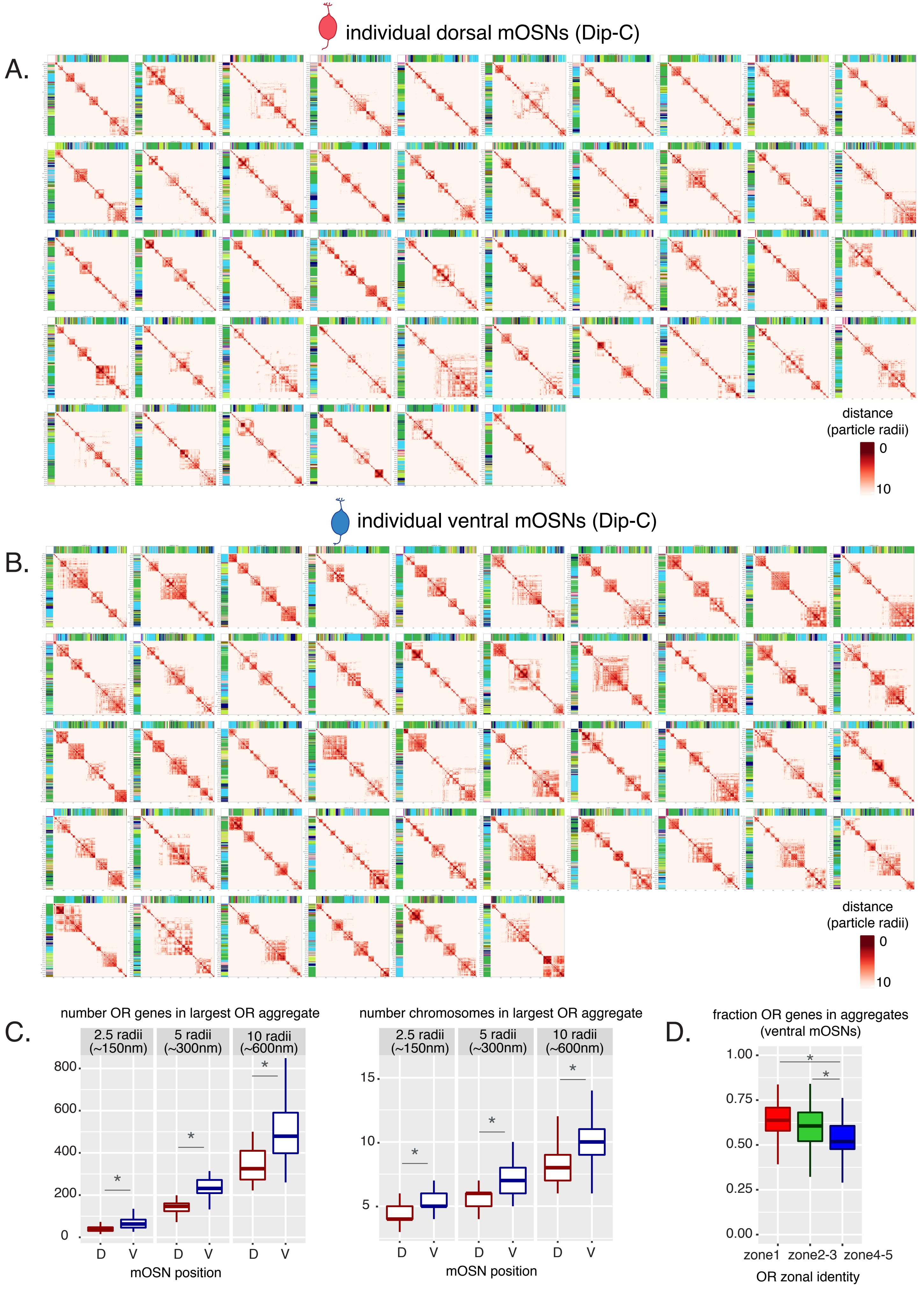

### Supplemental Figure 4

#### Supplementary Figure S4

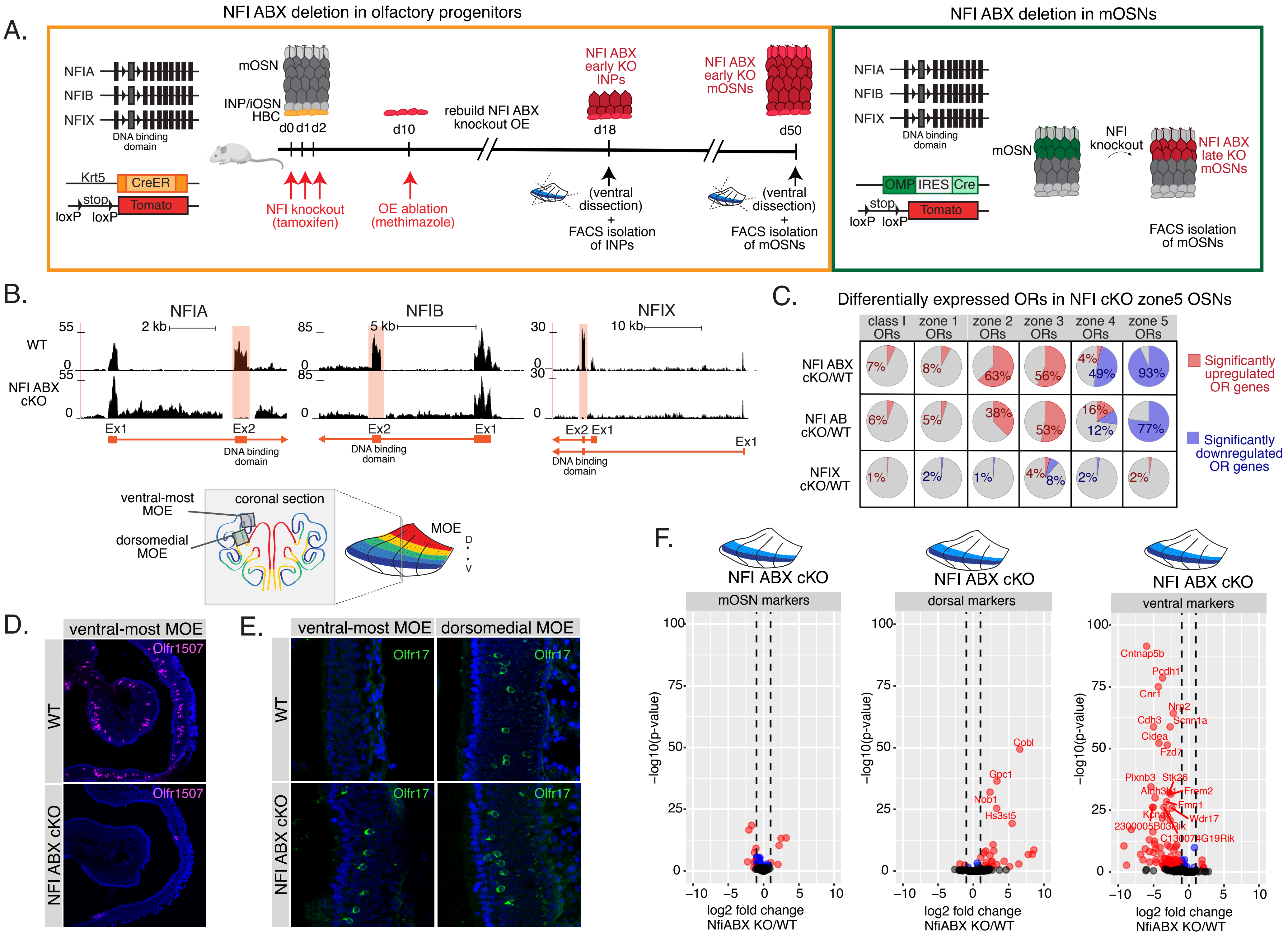

### Supplemental Figure 5

#### Supplementary Figure S5

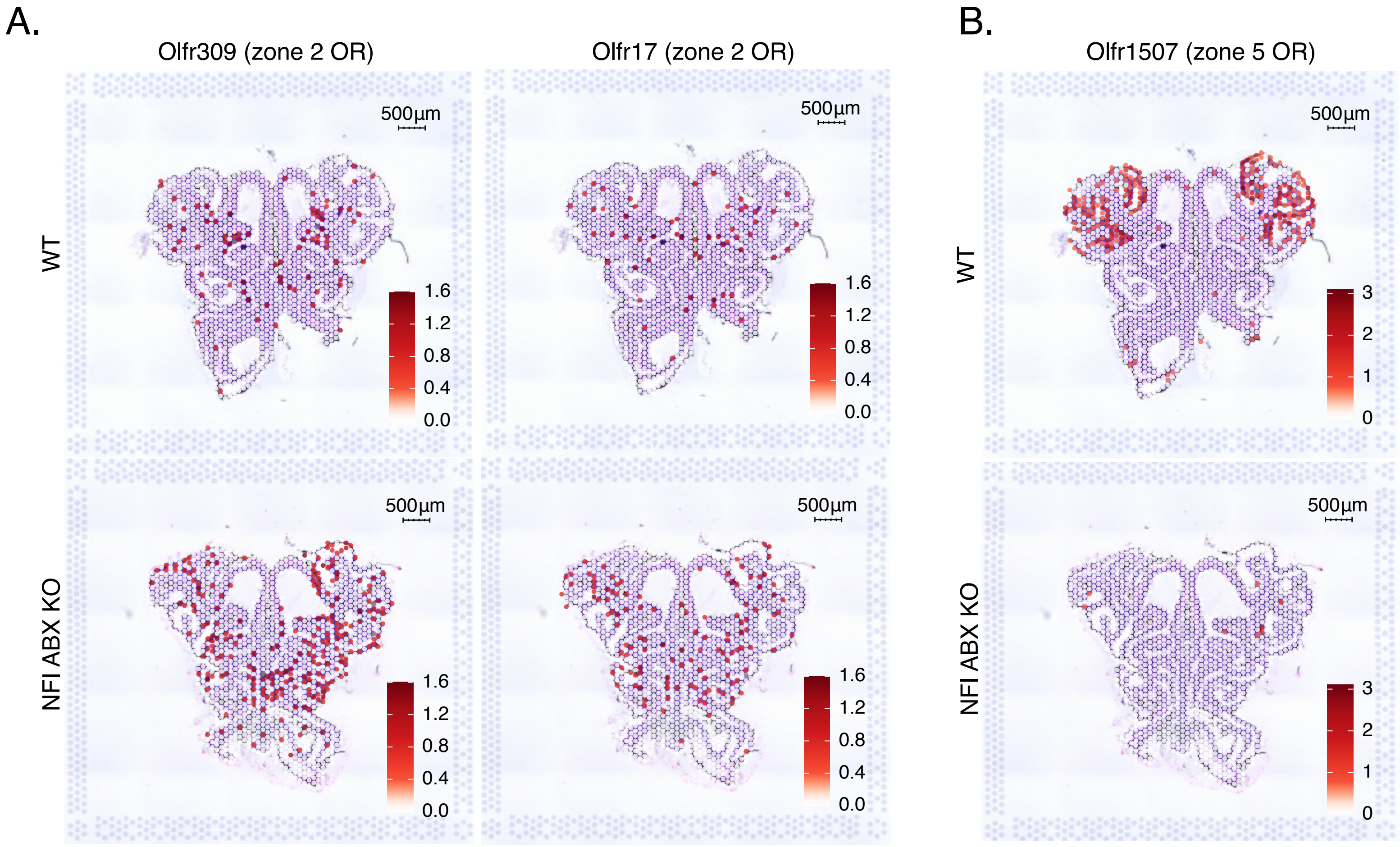

### Supplemental Figure 6

#### Supplementary Figure S6

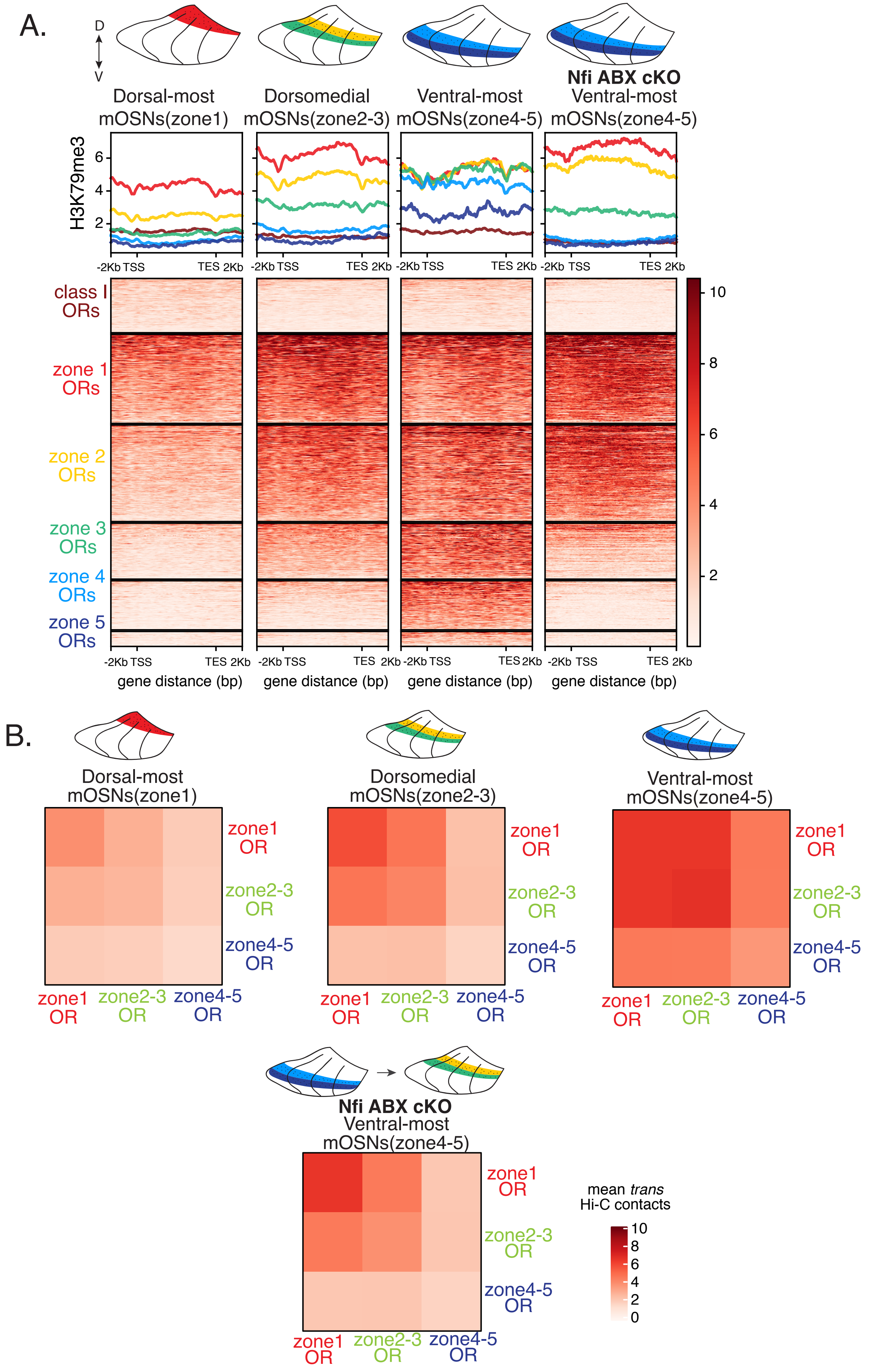

### Supplemental Figure 7

Supplementary Figure S7

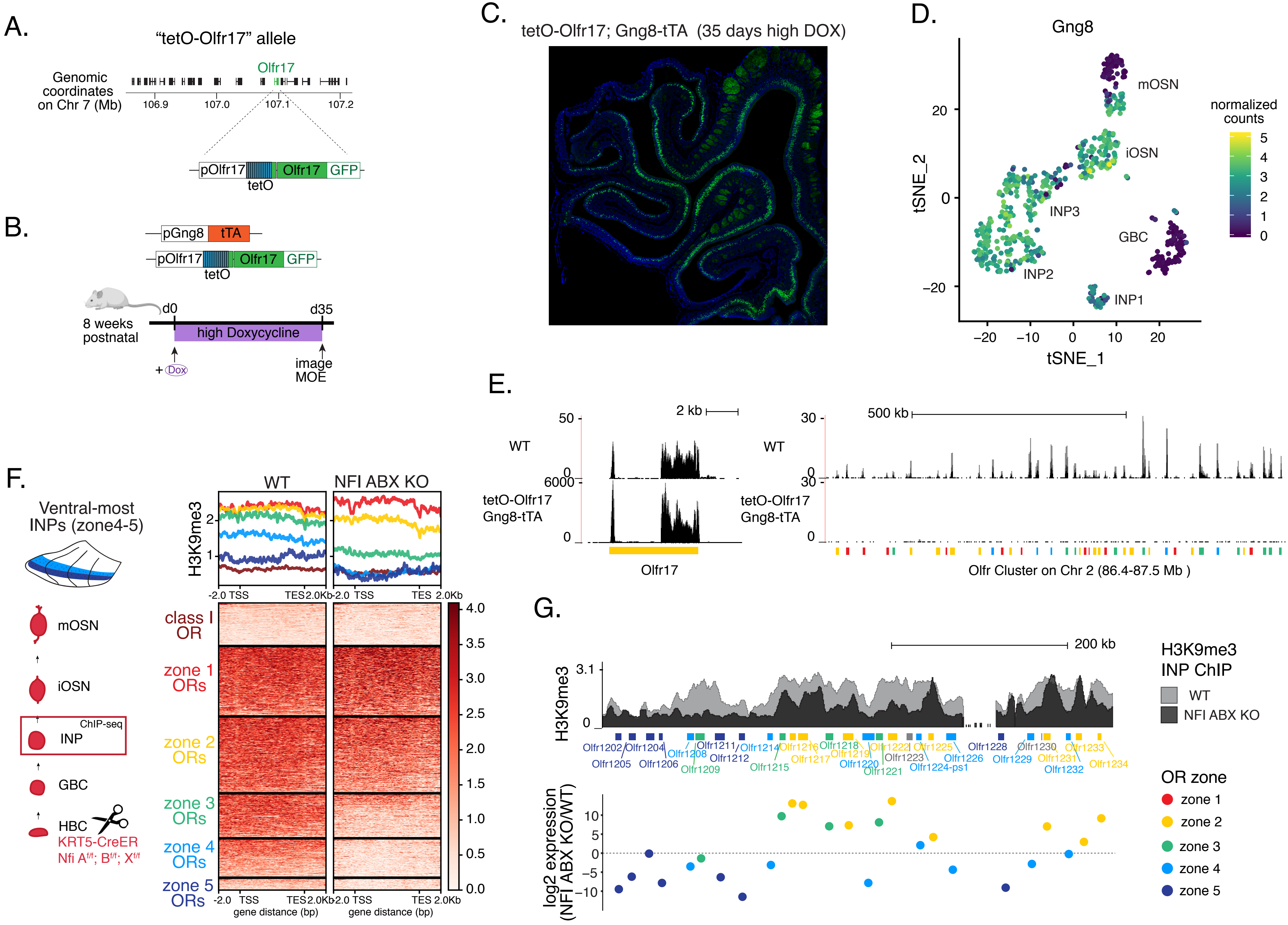
